# *Brachypodium* antifreeze protein gene products inhibit ice recrystallization, attenuate ice nucleation, and reduce immune response

**DOI:** 10.1101/2022.02.18.480559

**Authors:** Collin L. Juurakko, George C. diCenzo, Virginia K. Walker

## Abstract

Antifreeze proteins (AFPs) from the model crop, *Brachypodium distachyon*, allow freeze survival and attenuate pathogen-mediated ice nucleation. Intriguingly, each *Brachypodium* AFP gene encodes two proteins, an autonomous AFP and a leucine-rich repeat (LRR). We present structural models suggesting that ice-binding motifs on the ~13 kDa AFPs can “spoil” nucleating arrays on the ~120 kDa bacterial ice nucleating proteins that form ice at high sub-zero temperatures, consistent with decreases in ice nucleating activity by lysates from wildtype compared to transgenic *Brachypodium* lines. Strikingly, the expression of *Brachypodium* LRRs in transgenic *Arabidopsis* inhibited an immune response to pathogen flagella peptides (flg22), with structural modelling suggesting this was due to affinity of the LRR domains to flg22. Thus, these genes play distinctive roles in connecting freeze survival and anti-pathogenic systems via their encoded proteins’ ability to adsorb to ice as well as attenuating bacterial ice nucleation and the host immune response.

## Introduction

Since they cannot escape, plants must cope with a multitude of stresses, no more so than when winter approaches and they must defend themselves against ice formation, dehydration, and the pathogens that thrive even under snow cover. Do these multi-pronged assaults require the synthesis of specialised single-function defensive proteins or has evolution selected for multi-functional protectors? More than two decades ago, Marilyn Griffith argued the latter, in that certain hydrolytic proteins could have dual functions; they were antipathogenic and could also protect against ice-mediated damage^1,2^. Such proteins, however, did not have impressive antifreeze protein (AFP) activities thus her observations were not fully embraced by the ice-binding protein field. However, AFPs from a grass, the false brome model cereal *Brachypodium distachyon* (hereinafter, *Brachypodium*), derive from a putative bi-functional post-translational product, making them an obvious target for the investigation of abiotic stress resistance and antipathogenic multifunctionality.

*Brachypodium* AFPs (*Bd*AFPs) are encoded by a family of ice-recrystallization inhibition (IRI) genes, *BdIRI*s, encoding products that are exported to the apoplast and processed into two proteins: a leucine-rich repeat (LRR), derived from the amino-terminal domain, and a carboxy-terminal AFP^3,4^. Knockdown of the *BdIRI* translation products in transgenic *Brachypodium* as well as the heterologous expression of grass AFP in *Arabidopsis* have demonstrated that AFP activity confers protection against membrane electrolyte leakage subsequent to sub-zero temperature exposure^5,6^. Unlike AFPs from freeze-susceptible organisms, *Brachypodium* AFP (*Bd*AFP) does not lower the freezing point of solutions more than ~0.1 °C, but it is very effective at IRI, or the prevention of ice crystal coalescence at high sub-zero temperatures or under freeze-thaw conditions, a property crucially important for freeze-tolerant grasses. *Bd*AFP has been modelled based on its 67% identity to AFP from another grass *Lolium perenne* (*Lp*AFP) and has a beta-solenoid structure with two flat ice-binding motifs^5,7^. Indeed, although an order of magnitude smaller (~13 kDa), *Bd*AFP is structurally similar to the concatenated beta-solenoid repeats modelled for the ~120 kDa ice-nucleation proteins (INPs) produced by ice nucleation-active plant pathogens, such as *Pseudomonas syringae*. These bacteria can kill plants by inducing ice formation at high sub-zero temperatures in order to access intracellular nutrient stores and are of commercial concern since they are responsible for up to $7 billion in annual crop losses^8–11^. Notably, in addition to their IRI abilities, *Bd*AFPs appear to disrupt INP activity^12^, but structural uncertainty of the large INPs have hitherto thwarted speculation about the interactions of these two proteins.

Less is known about the LRR proteins derived from the *BdIRI*s. They have been modelled to the extracellular domain of the FLS2 receptor kinase of *Arabidopsis*^3,5^. FLS2 recognises the flg22 pathogen-associated molecular pattern (PAMP) derived from the flagellin of *P. syringae* and other bacteria, following which FLS2 triggers an immune response by initiating downstream biochemical changes and epigenetic reprogramming while energy is diverted from growth towards immune responses^13–15^. The similarity between the *BdIRI* LRR proteins and FLS2 suggested that they could bind flg22 PAMP and modulate the immune response. If so, *BdIRI*s could provide host protection against a costly immune response while the AFP domain neutralised one of *P. syringae’*s main weapons, INP. As noted, the structure and mechanisms of LRR:flg22 and AFP:INP interactions remain largely unknown.

The recent release of AlphaFold now provides a tool to investigate these relationships in concert with new experimental evidence on the anti-pathogenic properties of the two translation products.

## Results

### AFP-mediated attenuation of INP-induced freezing and INP models

The freezing point of *P. syringae* INP preparations was depressed by 1.55 °C with the addition of lysates from cold-acclimated (CA) wildtype *Brachypodium* compared to no lysate controls (*p* < 0.005, one way ANOVA; Figure 1; Table S1). No changes in ice nucleation temperature were seen with non-acclimated (NA) lysates, which have no measurable AFP activity. Likewise, nucleation temperatures of lysates from miRNA *BdIRI* knockdown lines (prOmiRBdIRI-1e or -3c^16^) with little to no AFP activity were not depressed, irrespective of their CA or NA treatment history, demonstrating that the attenuation of INP activity was dependent on AFP activity, as also shown by IRI assays (Figure 1).

**Figure 1.**
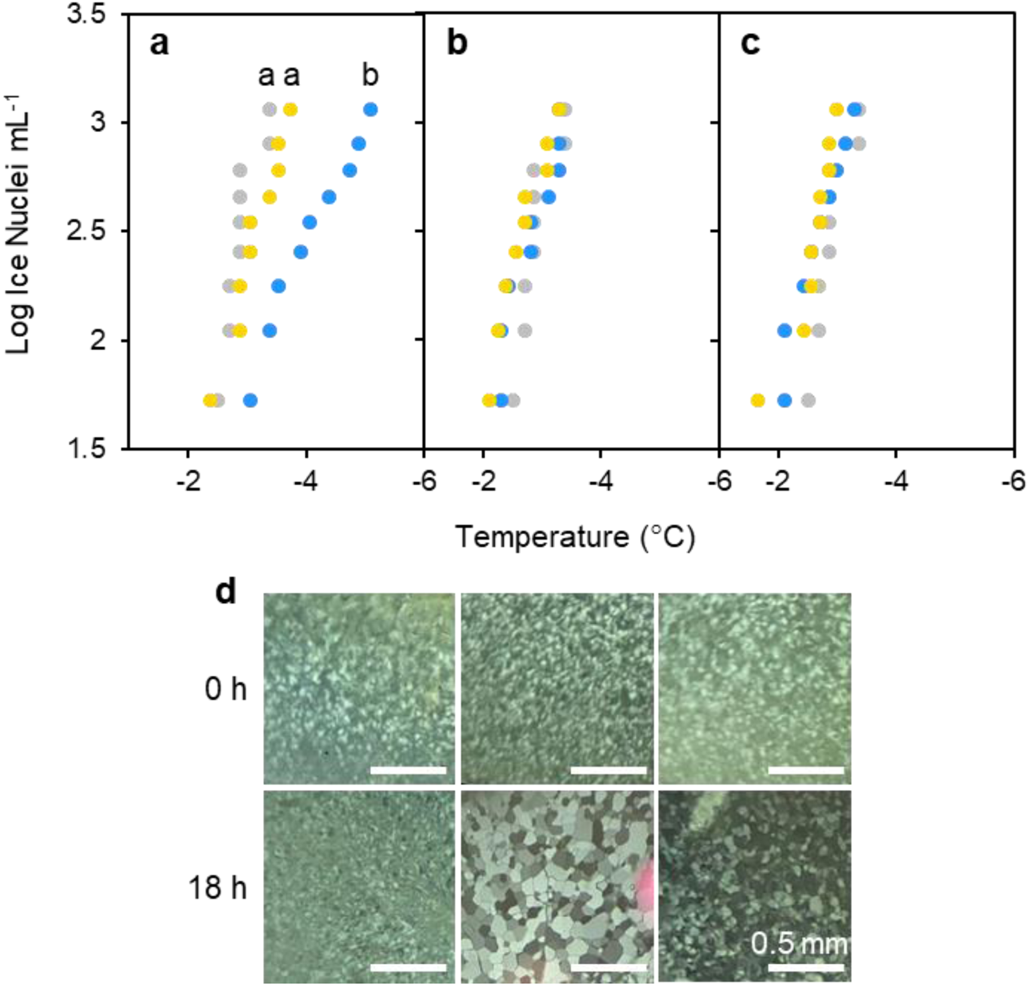
Impact of *Brachypodium* leaf tissue lysates or apoplast extracts on ice nucleation or ice crystal size. **a**, Representative ice nucleation assays conducted using ice nucleating protein (INP) preparations (50 µg mL^−1^) and concentrated leaf lysates of *Brachypodium* leaf tissue (final concentration of 20 mg mL^−1^). Small letter groupings indicating significance (*p* < 0.005, one way ANOVA) are shown as applicable. Samples (10 per plate) were repeated in triplicate with similar results. Comparison between INPs alone (grey dots) and INPs combined with lysates from cold-acclimated (blue) or non-acclimated (yellow) wildtype Bd21 lysates. Notably, the cold acclimated leaf lysates were comparably effective as the −1.26 °C depression of INP activity by purified recombinant *Bd*AFPs at equal concentration^12^. **b,** Comparison between INPs alone (grey dots) and INPs combined with lysates from the low temperature-induced transgenic *Bd*AFP knockdown line (prOmiRBdIRI-1e) that was either cold-acclimated (blue) or non-acclimated (yellow). **c,** Comparison between INPs alone (grey dots) and INPs combined with lysates from the low temperature-induced transgenic *Bd*AFP knockdown line (prOmiRBdIRI-3c) that was either cold acclimated (blue) or non-acclimated (yellow). **d,** Ice recrystallization inhibition activity in diluted (0.01 mg mL^−1^) apoplast extracts from leaves of cold-acclimated wildtype plants (left) and reduced activity in two cold-acclimated knockdown lines (prOmiRBdIRI-1e, middle, and prOmiRBdIRI-3c, right). Ice crystal sizes following an 18 h annealing period at −8 °C (lower images) are compared to those seen immediately after flash freezing (upper images). Scale bars represent 0.5 mm.

Attenuation of INP activity suggests a physical interaction between the two proteins. Accordingly, *in silico* modelling and docking predictions were performed using AlphaFold^17^ and FRODOCK^18,19^. Of the 7 isoforms, the product of *BdIRI1* was selected as the primary modelling representative, but all other full sequence models were made (Figure S1), as were just the LRR domains (Figure S2) or just the AFP domains (Figure S3). Previously, Phyre2 homology models of *Bd*AFPs alone were folded according to the crystal structure of *Lp*AFP^6,7^. The AlphaFold-generated model of the *BdIRI* primary translation product shows that the LRR and AFP domains are connected via a disordered linker that would facilitate endoprotease cleavage following secretion to the apoplast (Figure S1). Similar to the Phyre2 model, this new model folds each of the AFP domains into right-handed beta-solenoids consisting of opposing repetitive ice-binding a- and b-faces, with the sequence N**xVx**G/N**xVx**xG, where x is an outward facing, hydrophilic residue and the conserved triplets implicated in ice-association are indicated in bold (Figure 2a; Figure 2b; Figure S4a).

**Figure 2.**
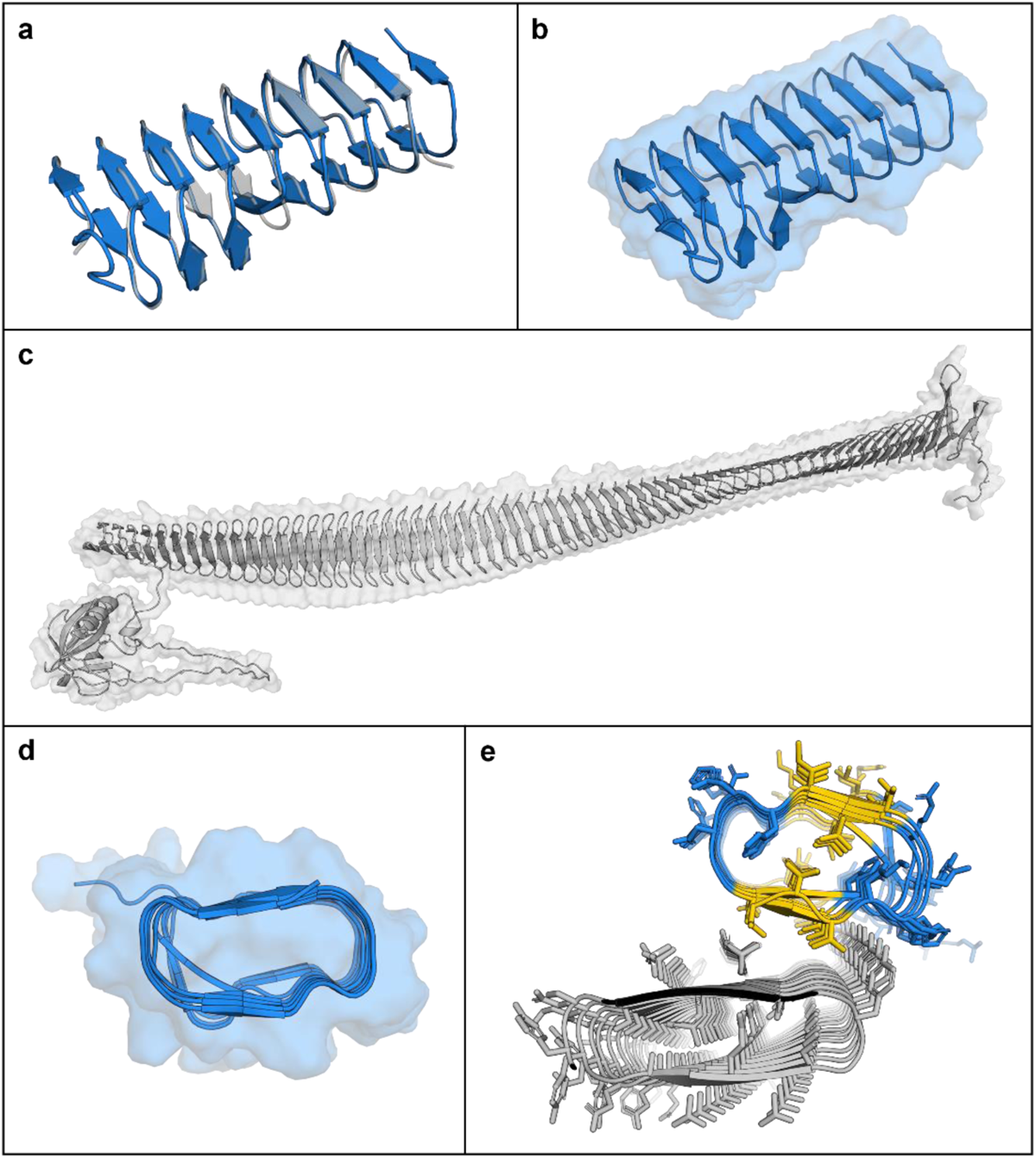
AlphaFold models of *Brachypodium distachyon* antifreeze protein (AFP) based on the representative *BdIRI1* AFP and *Pseudomonas syringae* ice nucleating protein (INP) and their interactions as predicted with FRODOCK. **a,** AlphaFold *Bd*AFP model (in blue) shown aligned to the crystal structure of the related *Lp*AFP (PDB ID: 3ULT; in grey). **b,** AlphaFold *Bd*AFP beta-solenoid fold with the solvent-accessible surface shown in transparency and with both ice binding surfaces depicted as flat ribbons (with the “a- and b-faces” on the upper and lower surfaces, respectively, with residues detailed in Figure S4). **c,** AlphaFold model of *P. syringae* INP as a twisted right-handed solenoid formed from the repetitive sequences with the N-terminal membrane anchor to the left, the cap sequence on the right, and the solvent-accessible surface area shown in transparency. **d,** *Bd*AFP cross section demonstrating the hydrophobic internal core and showing the relatively flat surface contours created by the ice-binding motifs, with the “a-face” shown on the upper side. **e,** Cross section of a FRODOCK docking prediction of interactions between *Bd*AFP (from *BdIRI1* AFP) in blue with the ice-binding faces in yellow (with the AFP “a-face”, closest to the INP), and *P. syringae* INP with two ice-binding faces (with the INP “a-face” upper, closest to the AFP). Note: The top 10 FRODOCK predictions (encompassing scores from 7212 down to 6349 as shown in Figure S5) show AFP binding along the entire length of the “a-face” of the INP via the “a-face” of the AFP, with only a representative image shown here.

Similarly, *P. syringae* INP (InaZ variant) was modelled as a representative INP. The bulk of the model shows a twisted ~28.5 x 0.25 nm beta-solenoid with dual, opposing flat ice-binding surfaces characterized by ~63 tandem <u>GYG</u>S**TQT**AxxxS**xLx**A repeats, where x is an outward facing hydrophilic residue, the presumptive water-ordering conserved triplets are shown in bold, and the putative interstrand dimerization tyrosine-ladder triplets are underlined (Figure 2c; Figure S4b). The opposing water-ordering surfaces appear to make equivalent contributions to INP activity^20,21^ and both appeared as flat a- and b-faces in the model. As shown, an N-terminal membrane anchor connects to the regularly ordered repeats of the beta-solenoid via a disordered linker composed of 61 residues (Figure 2c). The model also shows a cap-like structure at the carboxyl-end of the solenoid that terminates with a disordered tail of charged residues. The cap likely provides structural stability, preventing unwinding and amyloid fibril aggregation^22–24^.

Both AFP and INP models show that the conserved triplets of the water/ice-associating surfaces have two ranks of outward facing residues containing methyl groups, thought to be important for the arrangement of clathrate or ice-like water molecules^20^ (Figure S4). All of the top 10 scoring docking predictions show interactions through the a-faces of *Bd*AFP and INP (Figure 2d and e) with docking scores as high as 7200 (Figure S5). In contrast, the closely related *Lp*AFP, previously shown to attenuate INP less effectively than *Bd*AFP^12^, showed a lower maximum docking score of 4200, with 20% of models showing INP interactions on the single ice-binding a-face, and 80% on the opposite face (Figure S5; Figure S6a). A fish Type III AFP with little ability to perturb INP activity^6,27^, showed different possible interactions and still lower maximum scores of 2800 (Figure S5; Figure S6b).

Previous homology modelling of INPs based on the large bacterial repeats-in-toxin (RTX) proteins followed by manual inspection suggested that head-to-tail INP dimers could form through tyrosine ladders^28^. Here, FRODOCK and AlphaFold modelling applied to short INPs composed of 8 tandem repeats, necessitated by computational restrictions, also showed similar dimerization (Figure 3a). To experimentally test the INP model, ice nucleation assays were performed. Heating INP at 37 °C for 24 h depressed the freezing point (−3.33 °C), suggesting that proposed interstrand interactions may not fully reform again (Figure S7), consistent with the requirement for low temperature activation^29^. The polyphenol, tannic acid (TA), destabilises tyrosine ladders such as those found in amyloid fibres^30^, and thus INA assays were performed with TA as a further test of the modelled interstrand binding. As hypothesised, TA addition significantly depressed ice nucleation 2.28 °C more than INP preparations alone (*p* < 0.001, one way ANOVA; Figure 3b).

**Figure 3.**
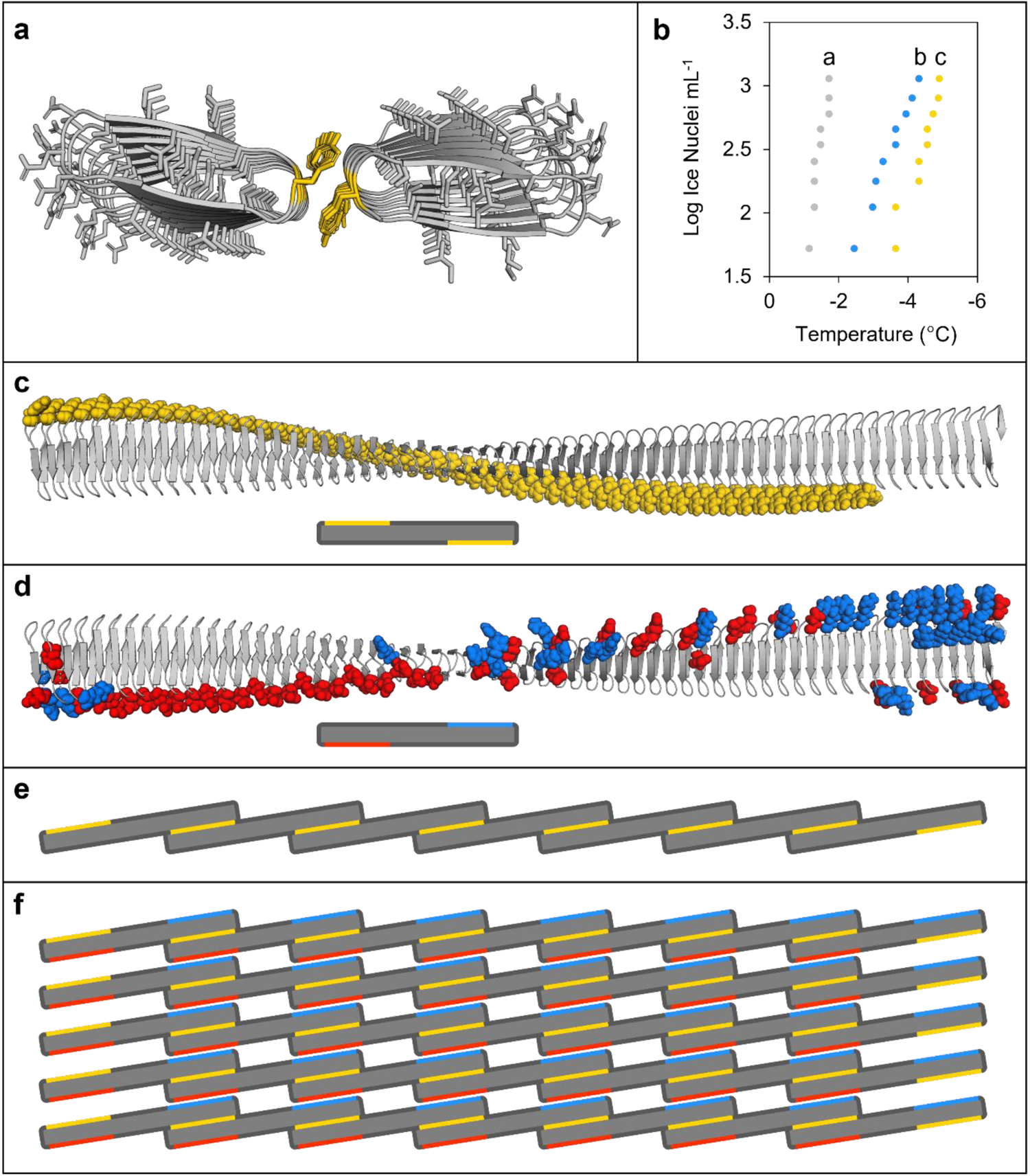
Modelling and other experimental evidence of INP interactions **a,** Tyrosine ladder interactions between INP monomers (InaZ) using FRODOCK and AlphaFold models with tyrosine residues in yellow showing the head-to-tail dimerization. **b,** Ice nucleation assay comparing ice nucleating protein (INP) preparations, at 50 µg mL^−1^ (grey dots) or when INPs were combined with 100 mM tannic acid (TA; blue dots) and INPs combined with 100 mM TA and *Bd*AFP from cold-acclimated wildtype lysates (yellow dots). Significance compared to INPs alone denoted by small letter displays with TA (*p* < 0.001, one way ANOVA) and TA + AFP (*p* < 0.001, one way ANOVA) and TA compared to TA + AFP (*p* < 0.01, one way ANOVA). Ice nucleation assays were repeated in triplicate with similar results. Note that the colligative depression of the TA solution was < 0.2 °C. **c,** An AlphaFold model of the *P. syringae* INP InaZ twisted beta-solenoid structure with GPI-anchor and linker hidden with tyrosine ladder residues highlighted in yellow (top) and a simplified illustration of the INP monomer with exposed tyrosine ladders represented by yellow bars (bottom). **d,** InaZ with charged residues on the opposing side of the beta-solenoid to the tyrosine ladder with negatively charged residues highlighted in red and positively charged residues in blue (top) and a simplified illustration with charged residues as described represented by bars (bottom). **e,** Schematic illustration of INP monomers (with the GPI anchor and cap sequence removed) organised by tyrosine ladder interactions, highlighted in yellow, to form a short INP filament. INP monomers would first partially dimerize with a single monomer forming tyrosine ladder interactions at each end of the twisted solenoid with two separate monomers on opposing sides. **f,** Self-assembly of INP filaments would precede the assembly of parallel filaments into aggregate sheets which could be stabilised through electrostatic interactions by the outward facing positively and negatively charged residues (red lines) found on opposing ends, respectively, on the side of the beta-solenoid opposite the tyrosine ladders (yellow lines). The schematic illustration shows 35 INP monomers (~250 nm^2^ of surface area) arranged into a sheet that would form patches on the surface of *P. syringae* (top down view). It should be noted that in theory, 34 INP monomers must interact to reach critical embryonic ice nucleus mass needed to achieve high sub zero ice nucleation at −2 °C.

Significantly, when *Bd*AFPs and TA were added together, the attenuation of INP activity was greater than with TA alone (−2.88 °C depression compared to control INP experiments; *p* < 0.001, one way ANOVA), suggesting that the “spoiling” of ice nucleation by TA and AFPs was likely via separate sites, again supporting our *Bd*AFP:INP interactive models. Based on models for the INP monomer (Figure 2c) and the modelled INP:INP interactions (Figure 3a), a representation of the INP oligomer filaments was made, stabilised by interactions between positively and negatively charged residues (Figure 3d, e). These were then used to form aggregate sheets (Figure 3f; Figure S8), consistent with the nucleation theory-predicted aggregation of 34 INPs required to mediate freezing at high sub-zero temperatures^31^.

### LRR-mediated attenuation of the host immune response and modelling

Representative LRR sequences corresponding to *BdIRI1*, *3*, and *7* were expressed in *Arabidopsis*, and the impact on the native immune response was assessed using oxidative burst assays (Figure 4). *Arabidopsis* lines expressing any of the LRRs showed significantly impaired responses (*p* < 0.001, one way ANOVA) when leaf disks were exposed to the immunogenic flg22-y peptide^32^, compared to wildtype *Arabidopsis* controls or transgenic plants bearing empty plasmids. Mean photon counts were reduced by 71% and 76% with LRR1, by 79% and 83% with LRR3 and by 64% and 70% with LRR7, when compared to transgenic and wildtype controls, respectively. No response to the flg22-ɑ peptide was seen in any of the samples, as expected given this peptide is non-immunogenic to *Arabidopsis*^32^.

**Figure 4.**
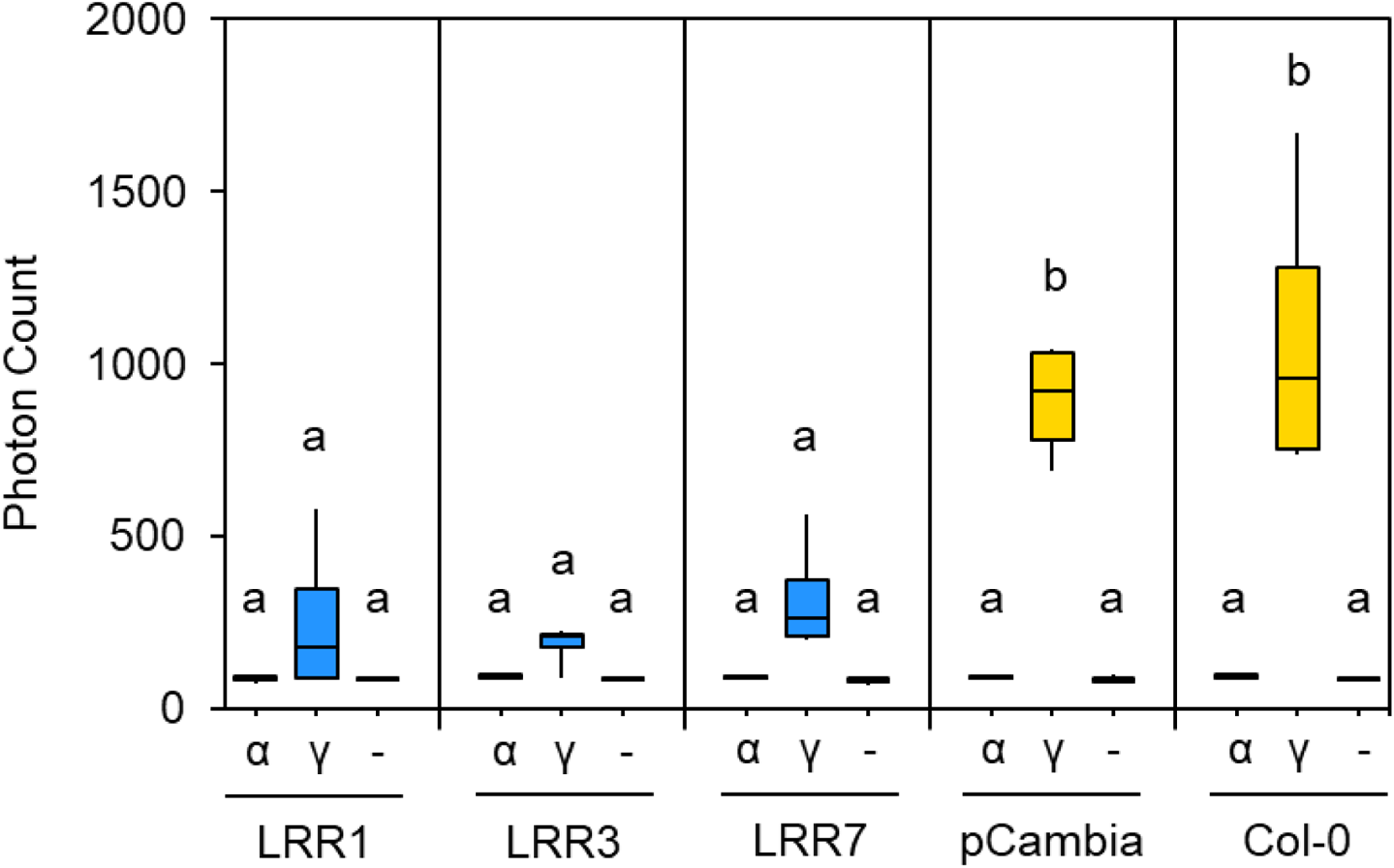
Representative oxidative burst assays (reactive oxygen species) as assessed by the measurement of emitted photons as a proxy for *Arabidopsis* immune response. Results are shown for untransformed wildtype *Arabidopsis* (Col-0), Col-0 plants transformed with an empty vector (pCambia1305.1, shown as pCambia), and *Arabidopsis* lines expressing the *BdIRI1, BdIRI3 and BdIRI4-*encoded LRR protein products shown as LRR1, LRR3, and LRR7, respectively. The flg22 epitopes used were the flg22-γ (shown as γ) and flg22-α (shown as α) peptides previously shown to be immunogenic and non-immunogenic, respectively, to *Arabidopsis*. No epitope blank controls (-) were also included. Small letter groupings represent statistically significant differences (*p* < 0.001, one way ANOVA) and assays were repeated in triplicate with similar results.

The AlphaFold LRR1 model showed similarity to the FLS2 crystal structure^33^ with opposing beta sheets and irregular alpha-helices forming a concave solenoid-like structure (Figure 5a). To further investigate experimental LRR:flg22 binding, AlphaFold interactions were modelled as described^34^, which predicted binding of the flg22-y epitope along the length of presumptive LRR binding pocket (Figure 5b-f).

**Figure 5.**
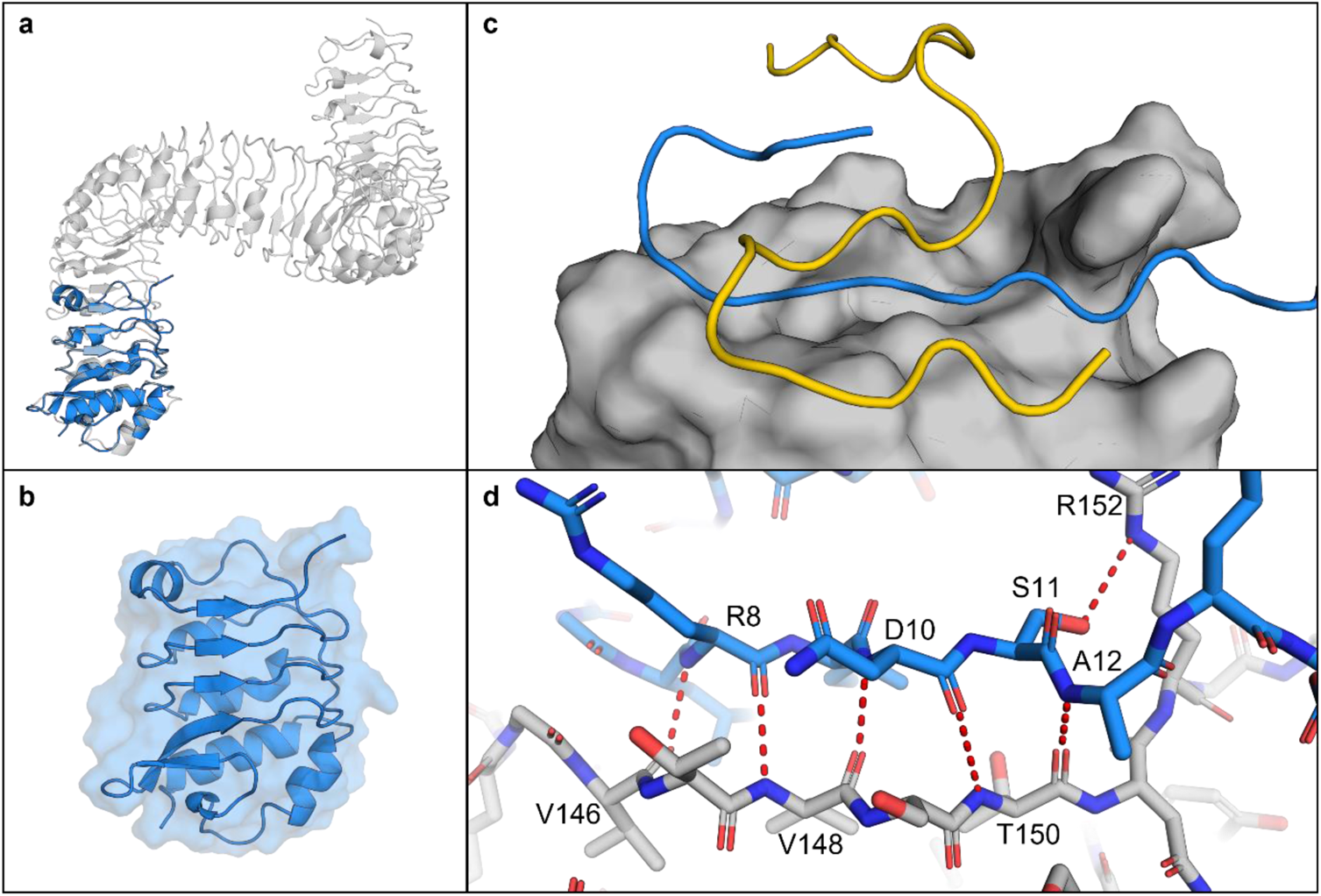
AlphaFold models of the *Brachypodium distachyon* LRR1 domain (based on *BdIRI1*). **a,** LRR1 in blue, aligned to the crystal structure of the *Arabidopsis thaliana* extracellular receptor FLS2 (PDB ID: 4MN8), in grey. **b,** LRR1 model showing secondary structure and solvent accessible surface area. **c,** LRR1 model with AlphaFold predicted binding of alpha (in yellow) and gamma (in blue) flg22 epitopes displaying a putative binding pocket on LRR1 with solvent accessible area shown. **d,** Hydrogen bonds between gamma residues (in blue) and LRR1 (in grey). Hydrogen bonds were predicted between Val-146, Val-148 and Arg-8; Val-148, Thr-150 and Asn-10; Thr-150 and Ala-12; and Arg-152 and Ser-11 with residues labelled. All hydrogen bonds were predicted using the PyMOL find polar contacts between chains function and are indicated by red dotted lines with lengths between 2.8 and 3.2 Å.

Five permutations of the flg22-y peptide sequence and modelling of these failed to show the same binding orientations and hydrogen bonds identified in the authentic LRR:flg22 complex, adding further support for the interaction between flg22 and *Brachypodium* LRRs (Figure S9). PDBePISA was used to further interrogate the interface between LRRs, including the representative LRR1 and the flg22-y or flg22-ɑ peptides, which predicted that binding occurs through a hydrophobic interface as indicated by a negative solvation free energy gain of −3.6 and −2.1 kcal mol^−1^, respectively (Table S2). The calculated interaction-specific surface areas for both peptides were substantive (*p* < 0.5; https://www.ebi.ac.uk/pdbe/pisa/), while interactions with permutations of the flg22-y sequence ranged from 10%-50% of the absolute value of the interface area between the LRR1:flg22-y complex (Table S2 and Figure S9). Using the same methodology, two LRRs together were also shown to have high interaction surface areas (Table S2; Figure 6). For example, when LRR3 and LRR4 were allowed to bind *in silico*, the heterodimer showed a negative solvation free energy gain of −7.3 kcal mol^−1^ when then bound to the flg22-y peptide, at least twice that of the LRR:flg22-y interaction, suggesting that more than one LRR could simultaneously bind to flg22-y *in planta*. Likewise, LRR3 and LRR4 homodimers showed negative solvation free energy gains of −9.1 kcal mol^−1^ and −11.9 kcal mol^−1^, respectively, with the LRR4 homodimer forming a sandwich around flg22-y stabilised by 25 predicted hydrogen bonds. By taking this idea further, modelled tetramers showed similar results (Figure 6g,h). Similar interactions were not seen with flg22-y permutation controls (not shown).

**Figure 6.**
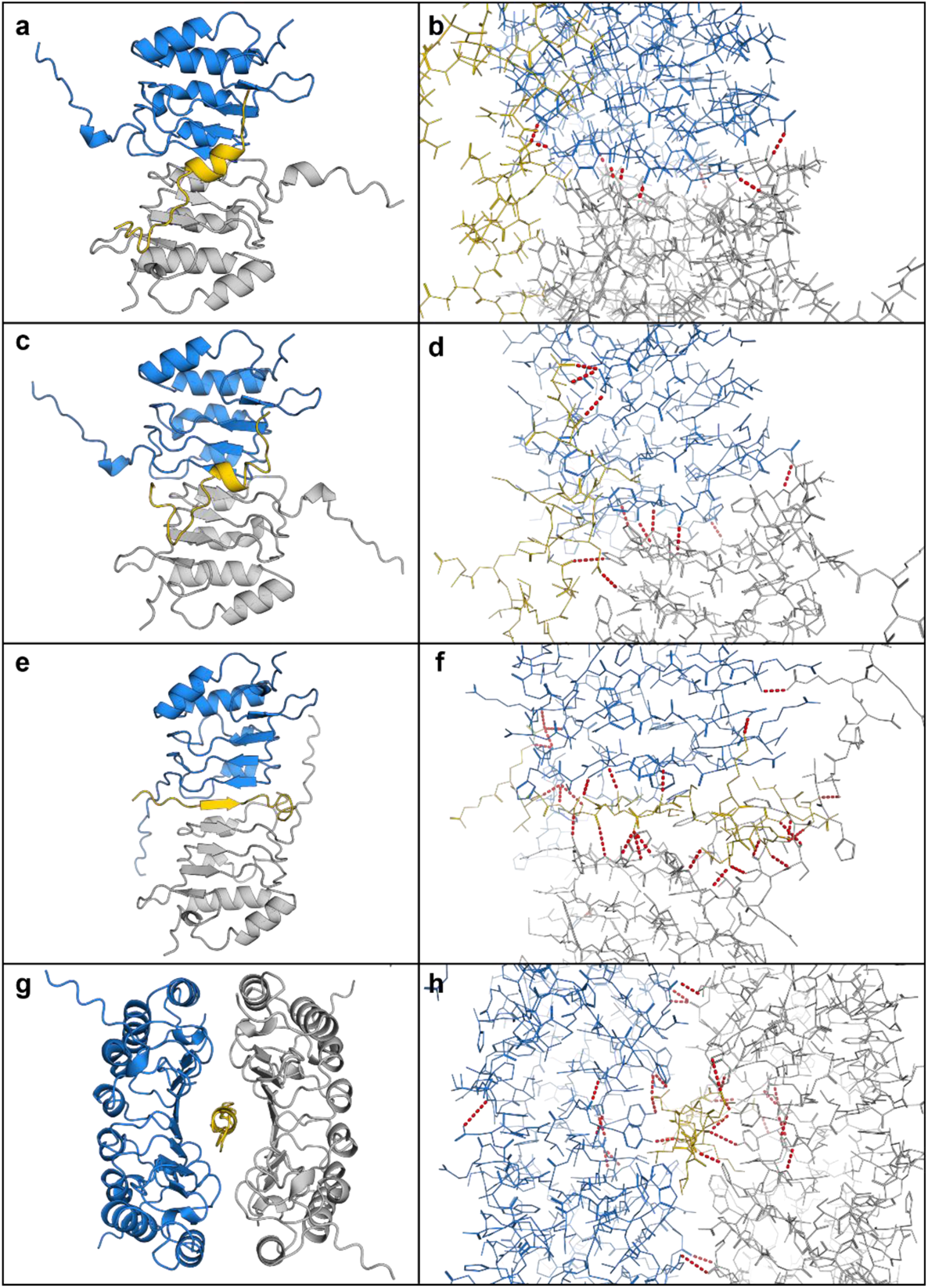
AlphaFold multimer heterodimer complexes with *Brachypodium* LRRs **a,** The LRR from *BdIRI3* (LRR3) in blue, *BdIRI4* (LRR4) in grey, and flg22-y in yellow. **b,** Hydrogen bonds in the LRR3 and LRR4 heterodimer and flg22-y complex. **c,** Two monomers of LRR from *BdIRI3* (LRR3) in blue and grey, and flg22-y in yellow. **d,** Hydrogen bonds in the LRR3 homodimer and flg22-y complex. **e,** Two monomers of LRR from *BdIRI4* (LRR4) in blue and grey, and flg22-y in yellow. **f,** Hydrogen bonds in the LRR4 homodimer and flg22-y complex. **g,** Two homodimers of LRR3, in blue and grey, for a total of four LRR3 monomers in complex with flg22-y, in yellow. **h,** Hydrogen bonds in the LRR3 homotetramer and flg22-y complex. All hydrogen bonds were predicted using the PyMOL find polar contacts between chains function and are indicated by red dotted lines with lengths between 2.8 and 3.2 Å.

## Discussion

The coming of winter is accompanied by low temperature-affiliated stresses in temperate plants^35^ including the orchestration of ice formation on *P. syringae’s* anchored INPs, presumably to freeze the apoplast at high sub-zero temperatures, allowing nutrient access for the pathogen. Such destructive activity must be neutralised by the plant, with ice formation allowed at lower temperatures, with additional safeguards in place to prevent ice crystal coalescence into larger membrane-damaging crystals. As well, light and water limitations would argue that energetically expensive immune responses must be muted. We now show that all of these requirements can be met by the *BdIRI* gene family. Thus, Griffith’s hypothesis that plants encode proteins that are both antipathogenic and ice protecting is correct^1^, but even she did not envision the remarkable attributes of the proteins encoded by *BdIRI*.

Investigations of how *Brachypodium* achieves these defences demand an insight into the physical interactions between proteins. However, INPs are a challenge due to their membrane association and their large ~120 kDa size. The previous TXT-homology folded INP model^28^ is largely concordant with our new AlphaFold model, with a twisted ~28.5 x 0.25 nm right-handed beta-solenoid repeat bearing flat putative water-organising a- and b-faces on opposite sides and with an amino-terminal cap and carboxyl terminal anchor domain (Figure 3). Similar to the homology-folded model, the new model combined with the FRODOCK docking algorithm on short INPs of 8 tandem repeats shows the potential for interstrand bonds through tyrosine ladder formation, which was tested by heating as well as TA addition, presumably targeting the cross-strand ladder^30^. Both of these treatments resulted in diminished ice nucleating activity, consistent with the known importance of dimerization for INP function, since by themselves, monomers don’t nucleate until −25 °C^36^. The tyrosine ladder shows an additional hydrophilic residue connection after energy minimization, suggesting that the tyrosyl hydroxyl group could function in conjunction with the flat faces (a-side; Figure S4), thus seamlessly extending the ice-organisation template. INP dimers must oligomerize to form INP filaments, and subsequently form aggregate sheets of aligned filaments that could be stabilised by electrostatic interactions of charged residues, also identified in the model. Such aggregates would minimally consist of the 34 INP monomers with a surface area of 240 nm^2^, covering ~10% of the length of the bacterium, sufficient to reach a critical embryonic ice nucleus mass at −2 °C^31^ (Figure 3).

How do *Bd*AFPs disrupt these vast INP water-organising templates, as demonstrated by the experimental attenuation of INP activity (Figure 1)? The docking models suggest that ranks of S/T residues on the a-face of *Bd*AFP dock near the ranks of the water-organising T on INP a-faces and possibly hydrogen bond, although this cannot be confidently predicted with the docking algorithm (Figure 3; Figure S4). The model further suggests that *Bd*AFP:INP interactions would not interfere with tyrosine ladder formation and the subsequent staircase architecture facilitating ice formation^26,35^. Nonetheless, since the docked *Bd*AFP appears close to the INP water-organising triplets, it is not surprising that the experimental freezing temperature was lowered. A distinct site is also supported by INP assays showing that the attenuation of INP activity is greater in the presence of both TA and *Bd*AFP than either alone. Substantive maximum docking scores bolster the model, with scores for *Bd*AFP:INP 2.6-fold higher than for INP:fish Type III AFP, which has more than 10-fold the thermal hysteresis antifreeze activity of *Bd*AFP^37^. Notably, fish AFP showing little or no inhibition of INP activity, supports the observation that attenuation of INPs is not an intrinsic property of AFPs, but rather an AFP-specific trait^6,27^. The grass *Lp*AFP with somewhat reduced ability to perturb INP activity showed two docking orientations involving the single *Lp*AFP ice binding face as well as the opposite flat face, in addition to docking scores between those shown by *Bd*AFP and fish AFP (Figure S5; Figure S8). When *Bd*AFP docks with the INP solenoids, we posit that ice formation initiates at slightly lower temperatures, reducing the probability that membranes surrounding the apoplast are breached, but then continues to function by protecting the plant against dangerous ice recrystallization with its two-sided ice binding faces, even as temperatures hover near 0 °C.

Although *Bd*AFP attenuates INP activity and thus eliminates a risk imposed by *P. syringae*, the presence of the bacterial flagella alerts the plant to a menace that typically results in the activation of the FLS2 receptor kinase through the LRR extracellular domain, leading to the mitogen-activated protein kinase cascade and signalling to direct energy to defensive proteins and away from growth and maintenance^15,38^. The *BdIRI* genes likely evolved from an ancestral rice phytosulfokine receptor tyrosine kinase with homology to the FLS2 immune receptor^3^. *P. syringae-*type flagellin proteins, flg22-y, interact with *Brachypodium* LRRs both experimentally and in the docking models, likely to attenuate immune responses (Figure 4; Figure 5). Notably, residues involved in flg22:LRR interactions are not the residues reported for flg22:FLS2^33^, suggesting an independent plant-specific mechanism. We speculate that the apoplast LRRs function as pattern receptor-like proteins, except that they lack an intracellular signalling domain. With their affinity for flagellin, these LRRs are hypothesised to bind to flagella from *P. syringae* and other pathogens and interfere with subsequent binding to FLS2, thus muting the *Brachypodium* immune response and its attendant energetic requirements.

It is possible that the LRRs work synergistically to attenuate the antimicrobial defences. LRR3 and LRR4, both expressed in leaves^5^, with a modelled disordered extension from the top of the “uncapped” solenoid and were predicted to self-assemble with 7 hydrogen bonds in a manner that aligned to FLS2 (Figure 6).

Indeed, analogous capless LRRs self-assembled^39^. The binding of flg22-y to the LRR dimer was tighter than to single LRRs and in a similar orientation with hydrogen bonding as shown in flg22-FLS2 binding (Table S2; Figure 6). As monomers or dimers or even tetramers, the LRR attenuates antipathogenic response in CA *Brachypodium*. The requirement for at least partial immune suppression is likely temporary. Once freezing takes place, the absence of free water may make pathogens less threatening and certainly, by the end of the CA period, the above-ground microbial community changes so that whereas *P. syringae* accounted for 8% of the taxa prior to transfer to 4 °C, 6 days later this taxon had disappeared^40^.

Taken together, the *BdIRI* gene products have demonstrated multiple functions that include anti-ice activity that partially “spoils” the ability of INP to form ice at very high sub-zero temperatures, strong IRI to protect plant membranes against freeze damage, as well as the attenuation of the plant immune response. These sequences present prime foci for the development of low temperature resilience in transgenic horticultural and agricultural crops. However, they also force us to re-evaluate an updated Beadle concept of one-gene, one-protein, one-function; here the protein products encoded by *BdIRI* have three distinct functions, an amazing “jab-cross-lead hook” combo to defy pathogens and freezing conditions alike.

## Methods

### Protein modelling and interaction predictions

INPs and AFPs were modelled using a version of AlphaFold (version 2.1.0) running on a Colab notebook using a high-RAM runtime provided by Colab Pro+^17^. Protein-protein docking predictions between various AFPs and INPs were performed using FRODOCK^18,19^. LRRs and flg22 binding predictions were modelled using AlphaFold with a 30 glycine linker connecting the amino terminal of the LRR to the flg22 peptide sequence as described^34^ with the LRR carboxyl apoplast localization signal sequences omitted to allow for more representative binding events. The LRR-linker-flg22 models were opened in PyMOL (version 2.4.1) and each was independently selected as a separate object and the linker was hidden. The LRR was then re-created as a closed surface to allow for binding pocket visibility. Any flg22 peptide secondary structures were hidden using the command “cartoon loop” and side chains were shown as sticks. Random permutations of flg22 were created using the^41^. Using the Protein Data Bank in Europe (PDBe) Protein, Interfaces, Structures, and Assemblies (PISA) tool (version 1.52), interfaces between LRRs and peptides were analysed and scored to quantify predicted binding affinity^42–44^. Chains were modified accordingly using the alter command and hydrogen bonds were predicted using the find polar contacts between chains command. The PyMOL commands used can be found in File S1. All sequences used for models can be found in File S2.

### Construction of LRR plasmids for *Arabidopsis*

Constructs of *BdIRI*s, synthesised by GeneART (Thermo Fisher Scientific, Waltham, MA, USA), were confirmed by Sanger sequencing (Platform de Sequencage, Laval, QC, Canada). Sequences containing LRR motifs confirmed by InterProScan^45^ were selected and used to form constructs in the pCambia1305.1 plant expression vector (Marker Gene Technologies Inc., Eugene, OR, USA).

Alignment and preparation of consensus reads from Sanger sequencing and alignment to reference sequences as well as *in silico* assembly of constructs was performed using Benchling (https://benchling.com/) (Benchling, 2020). Primers were designed using constructs containing 20 bp overlaps for Gibson assembly.

Amplicons of *Bd*LRRs containing overlaps were gel extracted using GeneJET Gel Extraction Kits (Thermo Fisher Scientific, Waltham, MA, USA) and pCambia1305.1 vectors were prepared and subsequently digested using NcoI restriction endonuclease prior to purification using Monarch DNA & PCR Cleanup Kits (New England Biolabs, Ipswich, MA, USA). Vector and insert concentrations were estimated using a Synergy H1 microplate reader (BioTek Instruments, Inc., Winooski, VT, USA) with a Take3 Micro-Volume Plate (BioTek Instruments, Inc.) and assembled using NEBuilder HiFi DNA Assembly Master Mix (New England Biolabs, Ipswich, MA, USA). Transformations were carried out using 2 µL of 4x diluted assembly mixes into DH5α cell lines, screened using colony PCR, and confirmed with Sanger sequencing. Positive clones were transformed into *Agrobacterium* AGL1 cell lines (Invitrogen, Carlsbad, CA, USA), and re-screened for the insert using colony PCR.

### Generation of transgenic *Arabidopsis*

Transformation for transient gene expression in *Arabidopsis* was performed as described^46^. *Arabidopsis* seeds were germinated and sown to potting soil and grown at standard conditions for three weeks. Simultaneously, *Agrobacterium* AGL1 cultures containing constructed plasmids were used to streak plates of YEB-induced agar containing appropriate antibiotics (Table S4^46^) and grown for two days before being scrapped and washed in 500 µL of wash buffer (10 mM MgCl2, 100 µM acetosyringone). Resuspended cells were gently vortexed and 100 µL was taken and diluted 10 times with infiltration buffer (25% MS, 1% sucrose, 100 µM acetosyringone, 0.01% Silwet L-77) with a final OD_600_ ≥ 12. Cells were then further diluted to an OD_600_ = 0.5. Finally, resuspended *Agrobacterium* cells were infiltrated into three-week-old *Arabidopsis* leaves on the abaxial side using a sterile 3 mL syringe with gentle pressure applied to the adaxial side. The zone of infiltration was marked gently using a black marker and labels were added to pots for transformant tracking. Following infiltration, plants were maintained under direct light for 1 h, using a consumer grade grow-op unit (SunBlaster Holdings ULC, Langley, BC, Canada), to allow for drying of the leaves to reduce drought response. Plants were then transferred to the dark for 24 h before returning to standard growth conditions in a climate-controlled chamber (see growth conditions below) for three days. Plants were removed from the chamber and leaf discs were taken using a sterilised 3.5 mm diameter biopsy tool (Robbins Instruments, Chatham, NJ, USA). Successful transformation and protein folding was confirmed using β-glucuronidase reporting screens on leaf discs from transformed leaves tandem to the ROS burst assay as described below.

### Plant material and growth conditions

*Brachypodium* seeds of an inbred line, ecotype Bd21 (RIKEN, Japan) and two transgenic AFP temporal knockdown lines^16^ were sown in potting soil and maintained at 4 °C in darkness for four days to synchronise germination. Seeds were moved to a climate-controlled growth chamber (Conviron GEN2000, Controlled Environments Ltd., Winnipeg, MB) at standard *Brachypodium* growth conditions of 70% relative humidity and 20 h days at ~150 μmol m^−2^s^−1^ at 24 °C followed by 4 h periods with no light at 18 °C. Plants were fertilised bi-weekly using 10-30-20 Plant-Prod MJ Bloom (Master Plant-Prod, Brampton, ON). Cold-acclimated (CA) plants were moved to a separate chamber (Econair GC-20, Ecological Chambers Inc., Winnipeg, MB) maintained at 4 °C and given a shortened day cycle of 6 h of light (~150 μmol m^−2^s^−1^) and 18 h dark for 48 h. Non-acclimated (NA) plants remained at standard conditions.

For *Arabidopsis* cultivation, wildtype seeds of ecotype Col-0 were soaked in water and placed at 4 °C in darkness for one week to synchronise germination before being sown in potting soil. Plants were grown in a climate-controlled growth chamber (Conviron GEN2000) under standard growth conditions of 16 h days at 22 °C with light at ~150 μmol m^−2^s^−1^ followed by 8 h of darkness at 20 °C. Prior to assay, plants were transferred to 15 °C for a day in the event that this facilitated proper folding of the LRR domains.

### Preparation of extracts and apoplast samples for AFP activity

For the analysis of AFP activity in wildtype and transgenic *Brachypodium*, extracts were prepared using a method modified from that previously described^47^. After acclimating three-week-old plants, 50 mg of leaf tissue was flash frozen with liquid nitrogen, ground into a fine powder, suspended in 400 µL of NPE buffer (25 mM Tris, 10 mM NaCl, pH 7.5, EDTA-free protease inhibitor tablets), and subsequently shaken for 4 h at 4 °C in the dark on a GyroMini nutating mixer (Labnet International Inc., Edison, NJ, USA). Samples were centrifuged at 13,000 × *g* for 5 min, chilled at 4 °C for 5 min, and the centrifugation and incubation were repeated.

The supernatant was transferred to 1.5 mL tubes and centrifuged again at 13,000 × g for 5 min before returning to 4 °C. Protein concentration was quantified using a Synergy H1 microplate reader (BioTek Instruments, Inc.) with a Take3 Micro-Volume Plate (BioTek Instruments) using A_280_ at a standard of 1 absorbance unit = 1 mg mL^−1^. Samples were normalised and diluted as described prior to assays, which included inspection of ice crystal morphology, electrolyte leakage and measurements of thermal hysteresis, all as described^5^. AFP assessment using IRI assays were done using plants prepared as indicated above^47^. Apoplast was prepared essentially as previously recommended^48^. After pipetting 10 µL of the apoplast samples 1 m onto a dry ice-chilled glass cover slip, they were annealed at −8 °C for 18 h. Images were captured through cross polarising film at 10x magnification, both before and at the end of the annealing period, with the experiment repeated independently at least three times for each sample.

### Ice nucleation assay

Assay of bacterial ice nucleation activity in the presence of AFPs and extracts was done as previously described^26,49^. Briefly, 2 µL of sample, containing either purified AFPs (1 mg mL^−1^), concentrated *Brachypodium* extracts (40 mg mL^−1^), or tannic acid (100 mM) combined with *P. syringae* INP (Ward’s Natural Science, Rochester, NY, USA) (at a final concentration of 50 µg/mL) or INP alone, was pipetted onto a polarised film in 10 replicates. After placing the film in a chamber, the temperature was lowered from −1 °C to −12 °C, at a rate of 0.2 °C min^−1^. Images along with the temperature were recorded every 60 s. The temperature at which 90% of the samples (T_90_) froze was considered to be the nucleation point. The logarithmic cumulative number of ice nuclei per mL in each sample (*K*(*T*)) was calculated as previously described^12^ using Vali’s equation^49^:

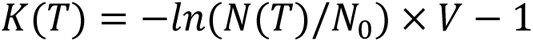

where *N*(*T*) represents the number of unfrozen samples remaining at time T, *N*_0_ represents the total number of samples, *V* represents the volume of the sample, and the *log*(*K*(*T*)) was taken to represent the logarithm of cumulative number of ice nuclei per mL. All 10 assays of each sample were repeated in triplicate.

### Immune response attenuation ROS burst assay

Plant immune response was assayed similar to that described^50^. Briefly, discs of transgenic *Arabidopsis* leaves were excised using a sterile 4 mm biopsy punch (Robbins Instruments, Chatham, NJ, USA) and Individually placed abaxial side down in sterile water (100 µL) in each wells of a white sterile 96-well microplate. Plates were incubated overnight at room temperature and then sealed with parafilm and covered with aluminium foil. After incubation, the water was carefully removed using a multichannel pipette, and replaced with 100 µL of solution containing 100 µM luminol, 10 µg mL^−1^ horseradish peroxidase (HRP), and 100 nM flagellin epitope ellicitor (or a control). Assays used a Synergy H1 Microplate Reader (BioTek Instruments, Inc.) using 1 s integration time, 60 min read time, with 2 min intervals measuring luminosity. Purified flg22 epitopes (EZBiolab, Parsippany, New Jersey, USA) included flg22-y (representative of the flg22 epitope of y-proteobacteria) and flg22-α (representative of the flg22 epitope of α-proteobacteria), which do and do not elicit an immune response in *Arabidopsis*, respectively^32^. The assays were conducted on extracts of wildtype Col-0, Col-0:*Bd*LRR1, Col-0:*Bd*LRR3, Col-0:*Bd*LRR7, and a Col-0:pCambia1305.1 empty vector control with both epitope treatments and a non-epitope blank controls. All assays were performed in triplicate.

### Statistical analysis

One way ANOVAs with post-hoc Tukey’s tests were performed in R using the package multcomp for compact letter displays of groups.

## Acknowledgments

We would like to acknowledge Dr. Robert L. Campbell for his generous modelling advice and for comments on the manuscript.

## Author Contributions

CLJ conducted all experiments and produced all figures. CLJ wrote the initial draft of the manuscript and all authors contributed to manuscript revision. GCD and VKW supervised the work.

## Supplementary Tables

**Table S1.** Ice nucleation assays with mean differences in T_90_ values (the temperature when 90% of samples nucleate) for ice nucleating proteins (INPs; 50 µg mL^−1^) in combination with cold-acclimated (CA) and non-acclimated (NA) wildtype Brachypodium distachyon lysates (Bd) and transgenic knockdown lines (prOmiRBdIRI-1e and prOmiRBdIRI-3c), or with tannic acid (TA; 100 mM), TA and CA-treated Bd, or with INPs pre-incubated at 37 °C for 24 h (heat) compared to INPs alone; in all cases values shown are the mean of three replicates with standard deviation. All experiments included buffer controls that did not nucleate (not shown).

| Sample | $\Delta T_{90}$ (°C) |
| --- | --- |
| INP + *Bd* (CA) | -1.55±0.28 |
| INP + *Bd* (NA) | 0.00±0.59 |
| INP + prOmiRBdIRI-1e (CA) | 0.52±0.32 |
| INP + prOmiRBdIRI-1e (NA) | -0.04±0.04 |
| INP + prOmiRBdIRI-3c (CA) | -0.03±0.37 |
| INP + prOmiRBdIRI-3c (NA) | 0.10±.24 |
| INP + tannic acid | -2.28±0.34 |
| INP + tannic acid + *Bd* | -2.88±0.54 |
| INP + Heat | -3.33±1.17 |

**Table S2.** Theoretical interface areas (Å^2^) from the PDBePISA interface output between representative receptor from the LRR translation product of BdIRI1 (LRR1), BdIRI3 (LRR3) and BdIRI4 (LRR4) and various flg22-ɑ or flg22-y ligand peptide chains, as well as the difference in total solvation energies of isolated and complexed structures where negative values indicate hydrophobic interfaces, or positive protein affinity, not including hydrogen bonds across interfaces (shown as Δ^i^G kcal mol^−1^), in addition to the observed solvation free energy gain where p < 0.5 indicates interfaces with higher than average hydrophobicity for the given structure suggesting the surface is interaction-specific (the Δ^i^G p-value).

| Protein (receptor) | Peptide (ligand)* | Interface Area ($\text{\AA}^2$) | $\Delta^iG$ kcal mol$^{-1}$ | $\Delta^iG$ $p$-value |
| --- | --- | --- | --- | --- |
| LRR1 | Gamma | 568.5 | -3.6 | 0.437 |
| LRR1 | Alpha | 232.8 | -2.1 | 0.415 |
| LRR1 | GammaP#1 | 115.1 | -0.3 | 0.600 |
| LRR1 | GammaP#2 | 54.6 | 0.0 | 0.544 |
| LRR1 | GammaP#3 | 168.5 | -0.6 | 0.415 |
| LRR1 | GammaP#4 | 291.4 | -0.8 | 0.441 |
| LRR1 | GammaP#5 | 198.8 | 0.1 | 0.433 |
| LRR3 | Gamma | 501.6 | -7.2 | 0.225 |
| LRR4 | Gamma | 467.3 | -6.8 | 0.243 |
| LRR3 # | LRR4 # | 1057.5 | -12.9 | 0.206 |
| LRR3 | LRR4 | 675.0 | -8.4 | 0.266 |
| LRR3+4 | Gamma | 567.9 | -7.3 | 0.207 |
| LRR3 # | LRR3 # | 853.4 | -12.4 | 0.161 |
| LRR3 | LRR3 | 588.1 | -8.8 | 0.211 |
| LRR3 x2 | Gamma | 757.1 | -9.1 | 0.286 |
| LRR4 # | LRR4 # | 674.1 | -6.5 | 0.401 |
| LRR4 | LRR4 | 887.7 | -7.4 | 0.371 |
| LRR4 x2 | Gamma | 1559.2 | -11.9 | 0.369 |
| LRR3 x4 | Gamma | 1224.7 | -6.7 | 0.516 |

\* GammaP# refer to random permutations of the flg22-γ peptide generated by the shuffle protein tool (https://www.bioinformatics.org/sms2/shuffle_protein.html).

## Supplemental Figures

**Figure S1.**
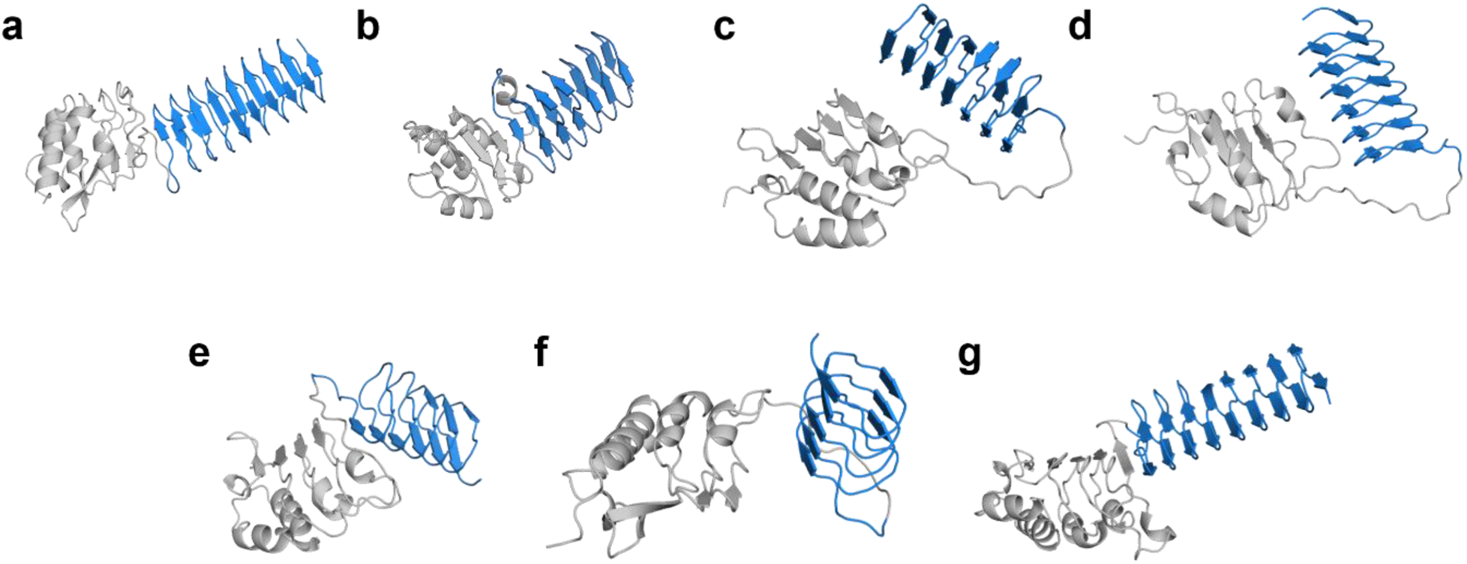
AlphaFold protein models of the 7 full *BdIRI* primary translation products with leucine rich repeat (LRR) domains in grey and antifreeze protein (AFP) domains in blue; the hydrolytic cleavage site is located between the two. **a,** *BdIRI1.* **b,** *BdIRI2*. **c,** *BdIRI3*. **d,** *BdIRI4*. **e,** *BdIRI5*. **f,** *Bd*IRI6. **g,** *Bd*IRI7.

**Figure S2.**
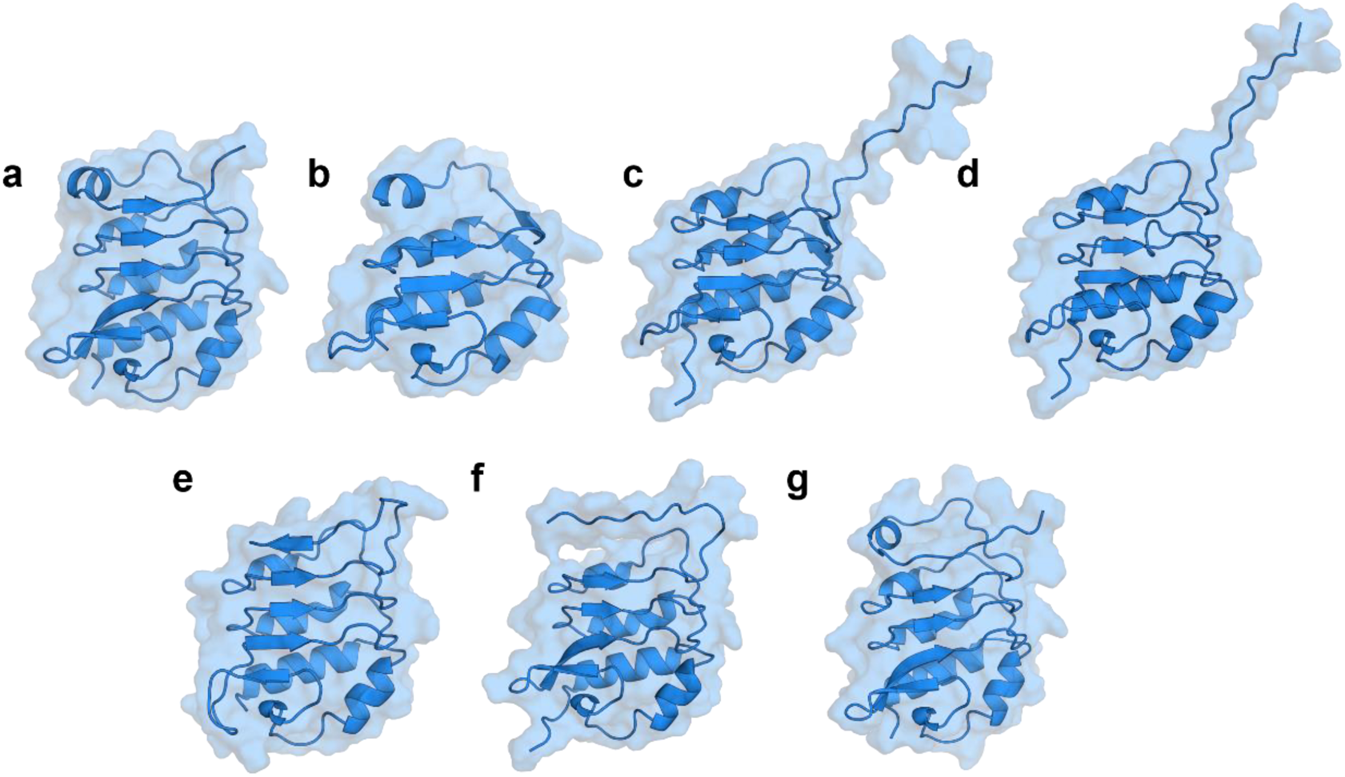
AlphaFold models of *BdIRI* leucine rich repeat (LRR) domains folded independently and showing the solvent-accessible surface area. **a,** LRR1. **b,** LRR2. **c,** LRR3. **d,** LRR4. **e,** LRR5. **f,** LRR6. **g,** LRR7.

**Figure S3.**
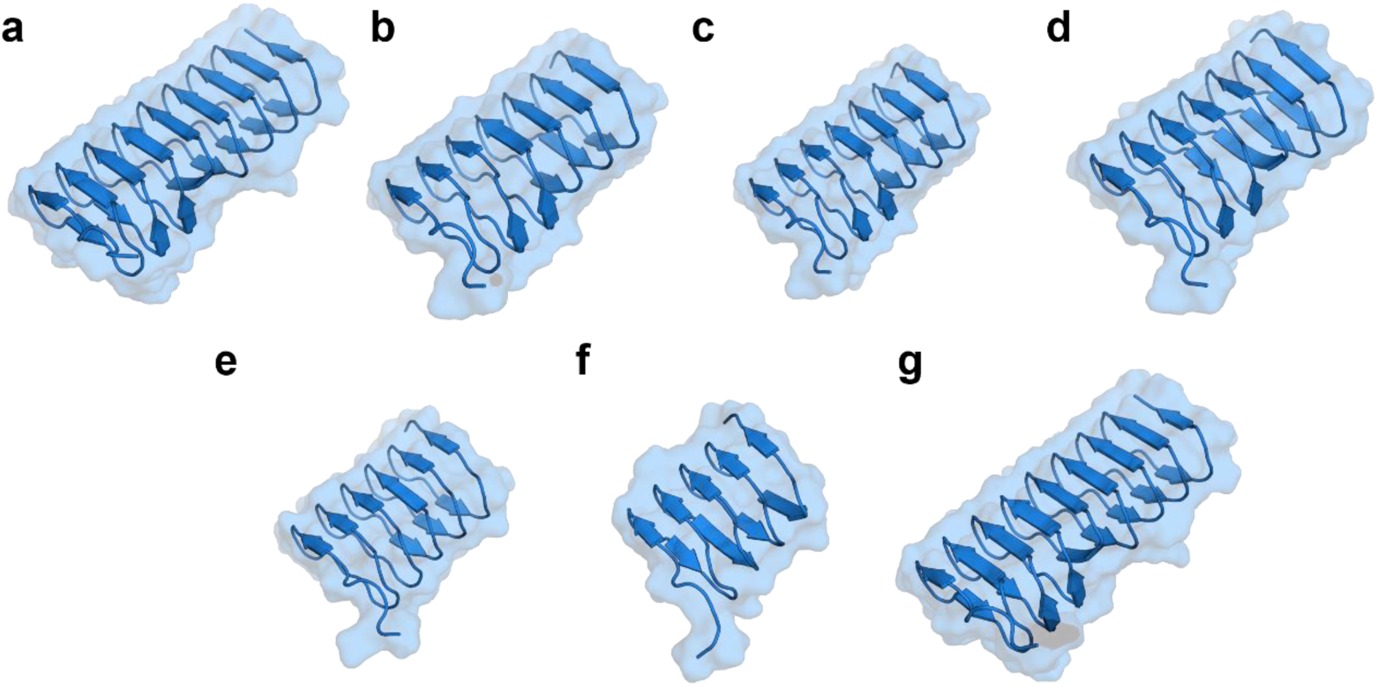
AlphaFold models of *BdIRI* gene antifreeze protein (AFP) domains folded independently and showing the solvent-accessible surface area. **a,** *Bd*AFP1. **b,** *Bd*AFP2. **c,** *Bd*AFP3. **d,** *Bd*AFP4. **e,** *Bd*AFP5. **f,** *Bd*AFP6. **g,** *Bd*AFP7.

**Figure S4.**
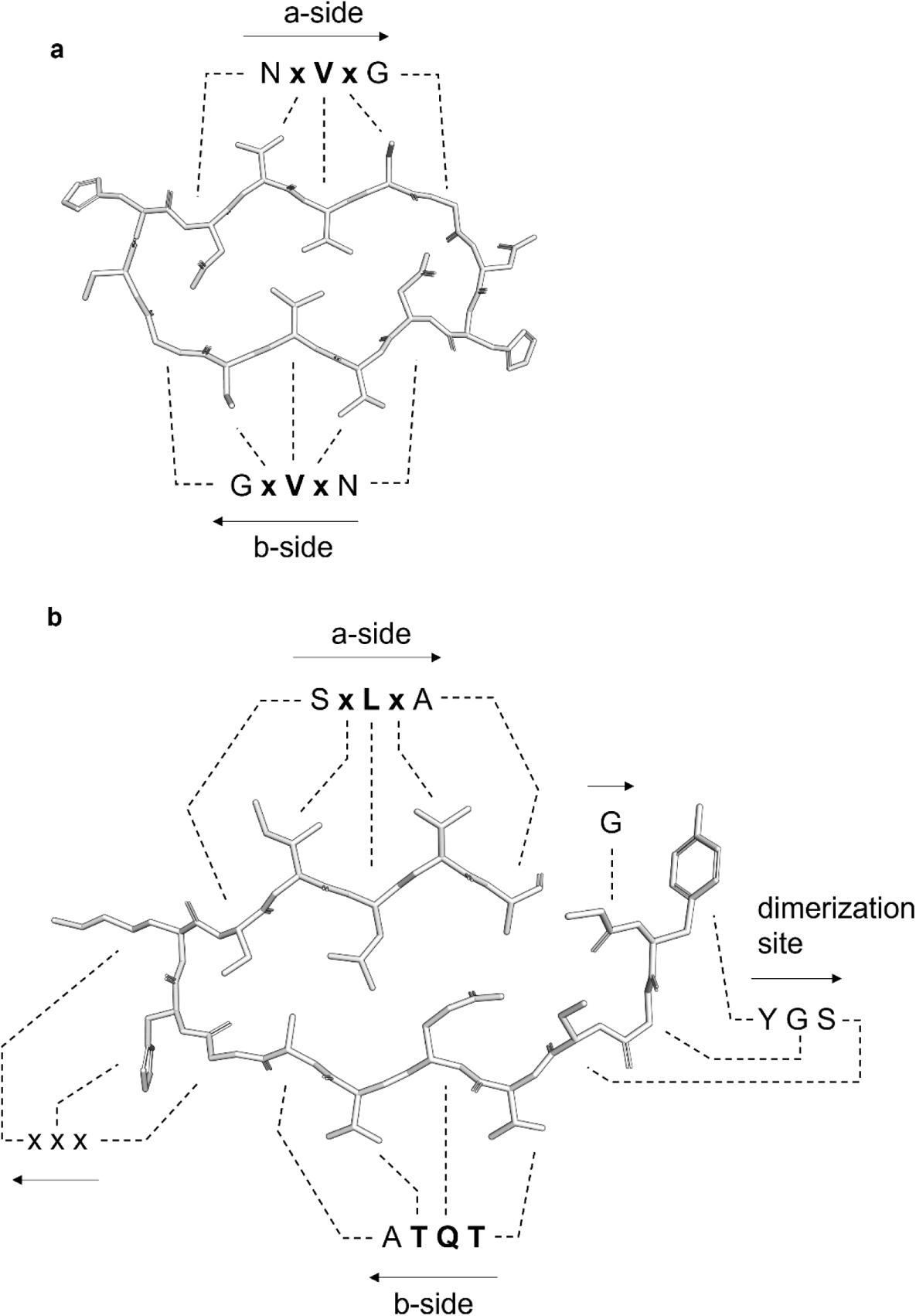
Stick representations of ice-binding motif repeats of AFPs and INPs. **a,** *Brachypodium distachyon* AFP repetitive “a and b ice-binding faces” on the top and bottom, respectively, of the cross-sectional depiction of the sequence N**xVx**G, where x is an outward facing, hydrophilic residue and ice-binding conserved triplets are indicated in bold. **b,** A single representative *Pseudomonas syringae* INP ice-binding tandem <u>GYG</u>S**TQT**AxxxS**xLx**A repeat, where x is an outward facing hydrophilic residue and the “a and b ice binding faces” on the top and bottom, respectively, are shown in bold, with the conserved sequence forming the tyrosine ladder also shown. Residues are labelled with dashed lines and arrows indicate amino- to carboxyl-sequence directionality corresponding to the model.

**Figure S5.**
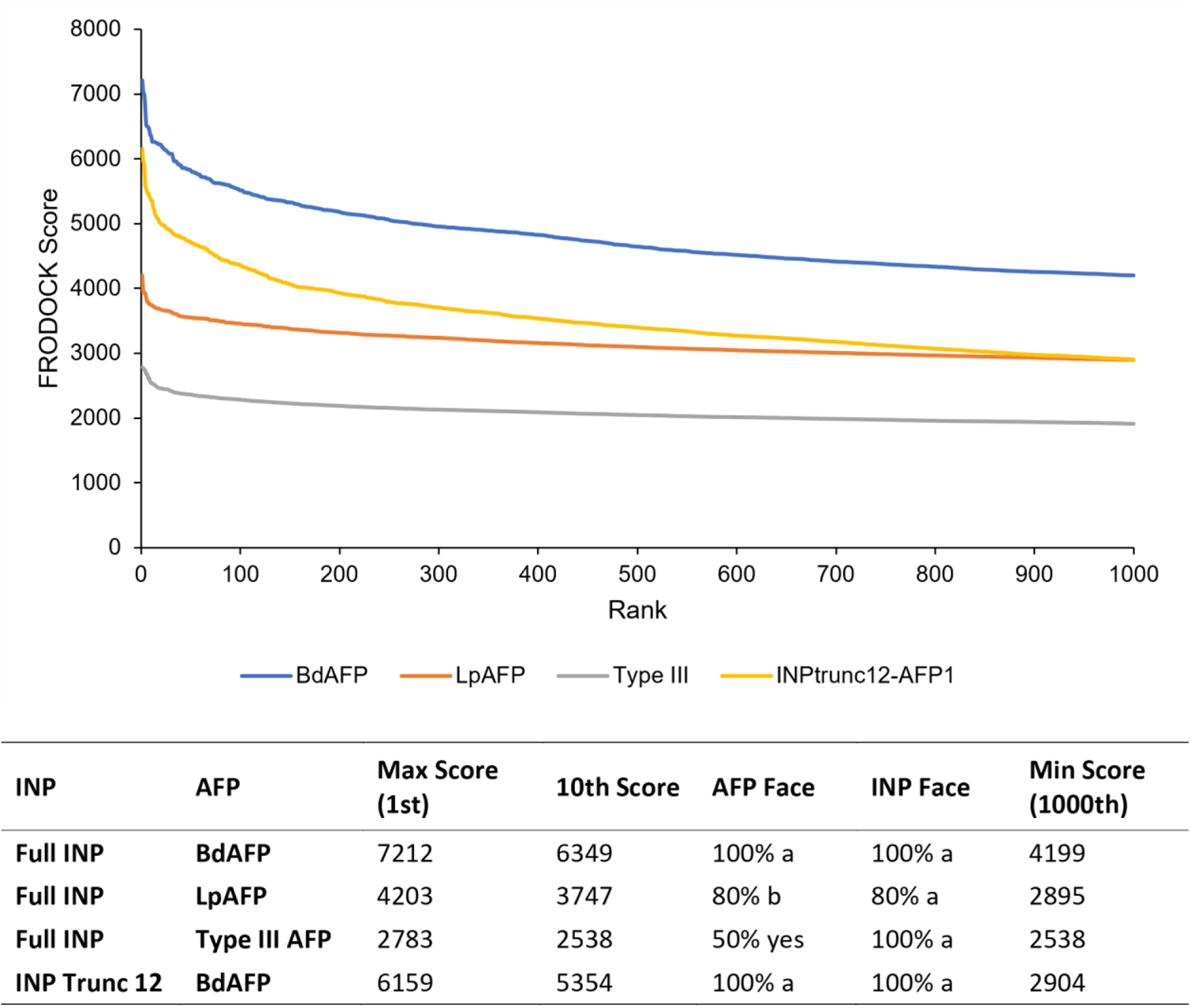
Summarised FRODOCK correlation scores and rankings for predicted docking of *Brachypodium distachyon* antifreeze protein (*Bd*AFP), *Lolium perenne* AFP (*Lp*AFP), and a Type III AFP from the fish, *Macrozoarces americanus*, against *Pseudomonas syringae* ice nucleating protein, INP, using the InaZ variant as a model as well as truncated INPs using a segment of the beta-solenoid containing 12 of the tandem repeats modelled with *Bd*AFP. The top 1000 rankings and scores are shown (top) and the max, 10th, and min (1000th) scores summarised and the INP and AFP faces predicted to interact are displayed based off the top 10 docking arrangements (below).

**Figure S6.**
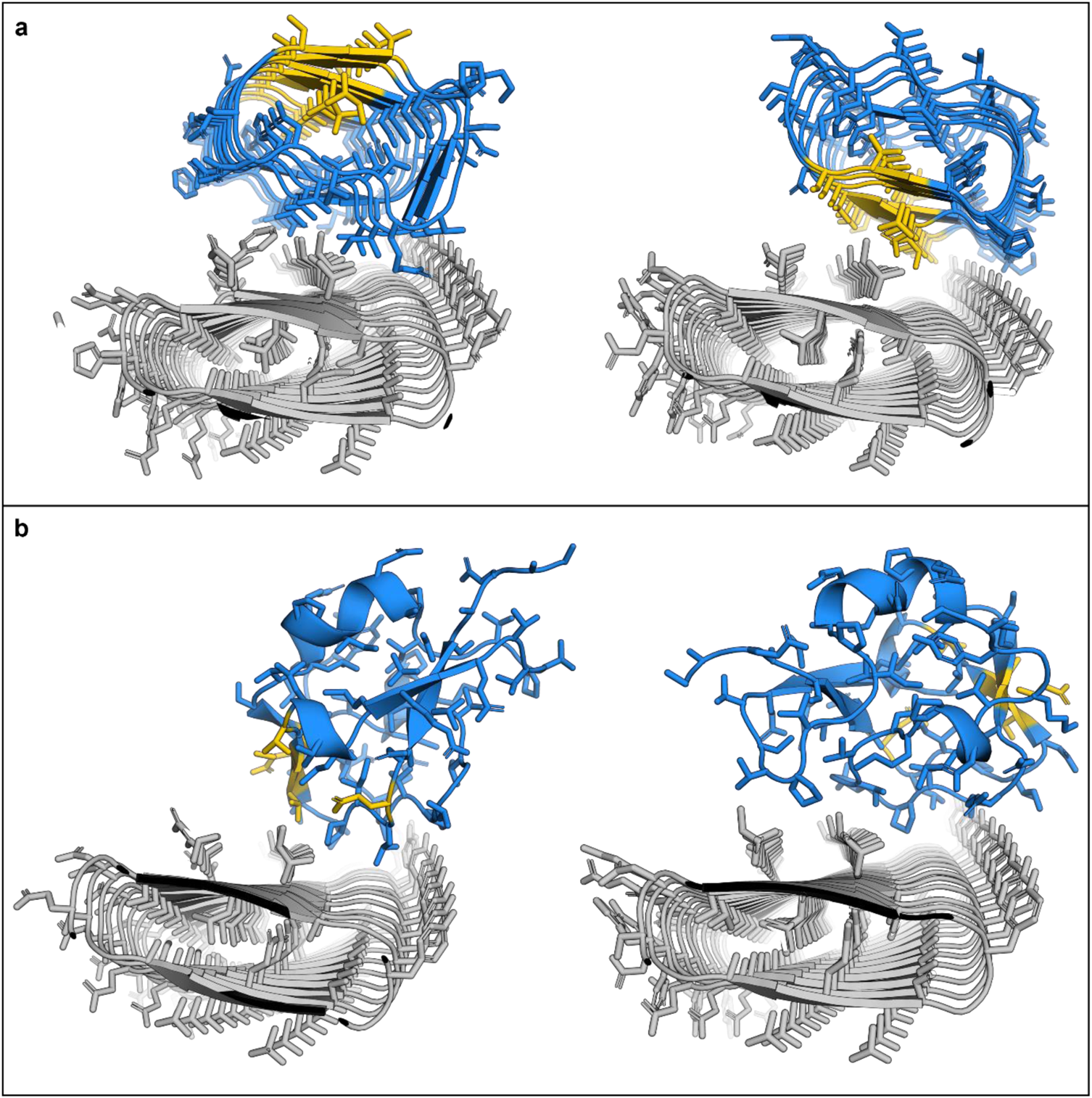
Cross sections of FRODOCK docking predictions of antifreeze proteins (AFPs) in blue with their ice binding residues in yellow, on the AlphaFold model of the *Pseudomonas syringae* ice nucleating protein (INP) in grey. Note that AFPs were predicted to bind along the entire length of the INP solenoid and only the top representative images shown. **a,** *Lolium perenne* AFP, *Lp*AFP, crystal structure model (PDB ID: 3ULT). Only the “a-side” of the AFP has ice-binding affinity^25^. The FRODOCK algorithm suggests that docking could occur on either face, thus two images shown involving the b-face (left as seen in 80% of the top models) or the opposite ice binding a-face (right) seen in 20% of the models. **b,** Type III AFP from *Macrozoarces americanus* crystal structure model (PDB ID:1MSI). Shown are two representative images, one showing docking that involves the ice binding face (left, seen 50% of the time) and the other elsewhere (right). Type III AFP does not attenuate bacterial ice nucleation^26^ and the docking scores are lower than those shown by *Bd*AFP or *Lp*AFP as stated in Results with details provided in Figure S5).

**Figure S7.**
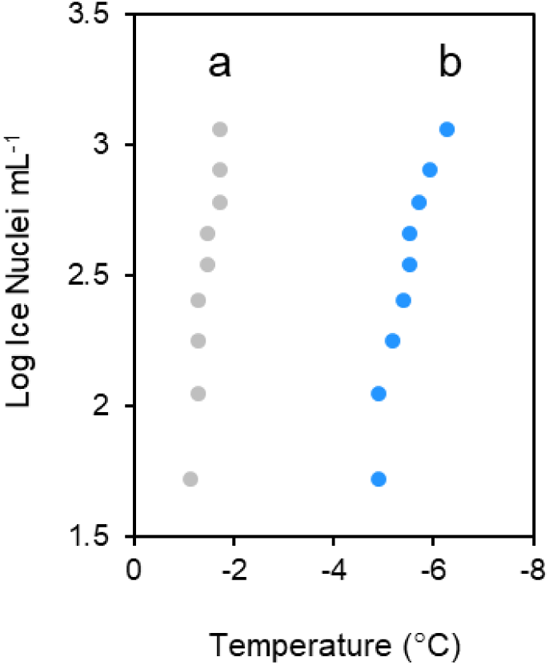
Ice nucleation assay comparing *P. syringae* ice nucleating protein (INP) preparations alone (in grey) to INPs maintained at 37 °C for 24 h (in blue) before assessment. Assay was repeated in triplicate with similar results. Small letter groupings indicating significance (*p* < 0.001, one way ANOVA) are shown. Samples (10 per plate) were repeated in triplicate with similar results.

**Figure S8.**
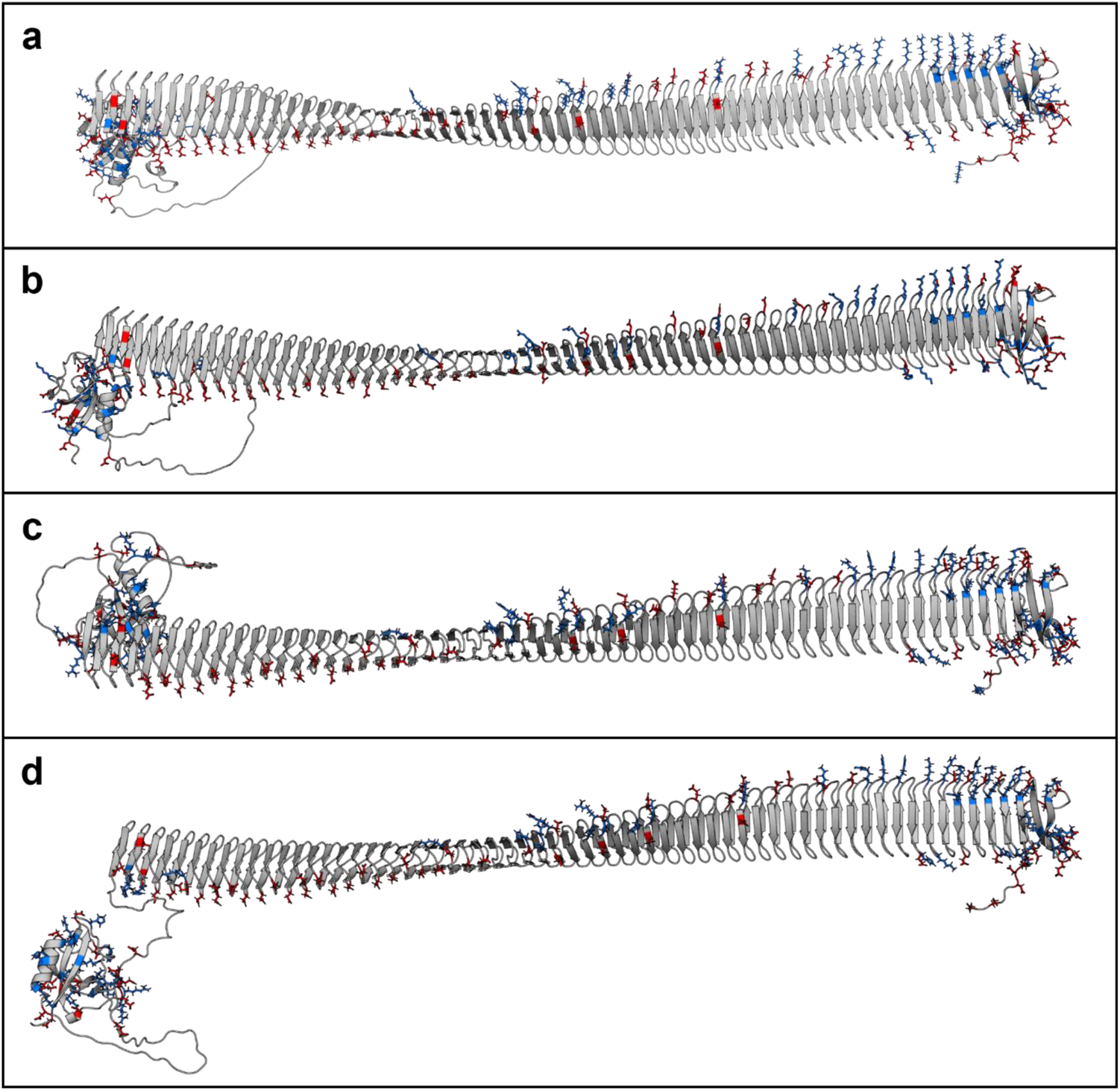
AlphaFold models highlighting charged residues of *Pseudomonas syringae* ice nucleating protein (INP) variants. **a,** The INP used as the model for this work, InaZ. **b,** variant InaV. **c,** variant InaK. **d,** variant InaQ. Negatively charged residues are highlighted in red and positively charged residues in blue. We propose that these positively and negatively charged residues may be involved in stabilising the formation of INP aggregate sheets by INP filaments formed through tyrosine ladder interactions.

**Figure S9.**
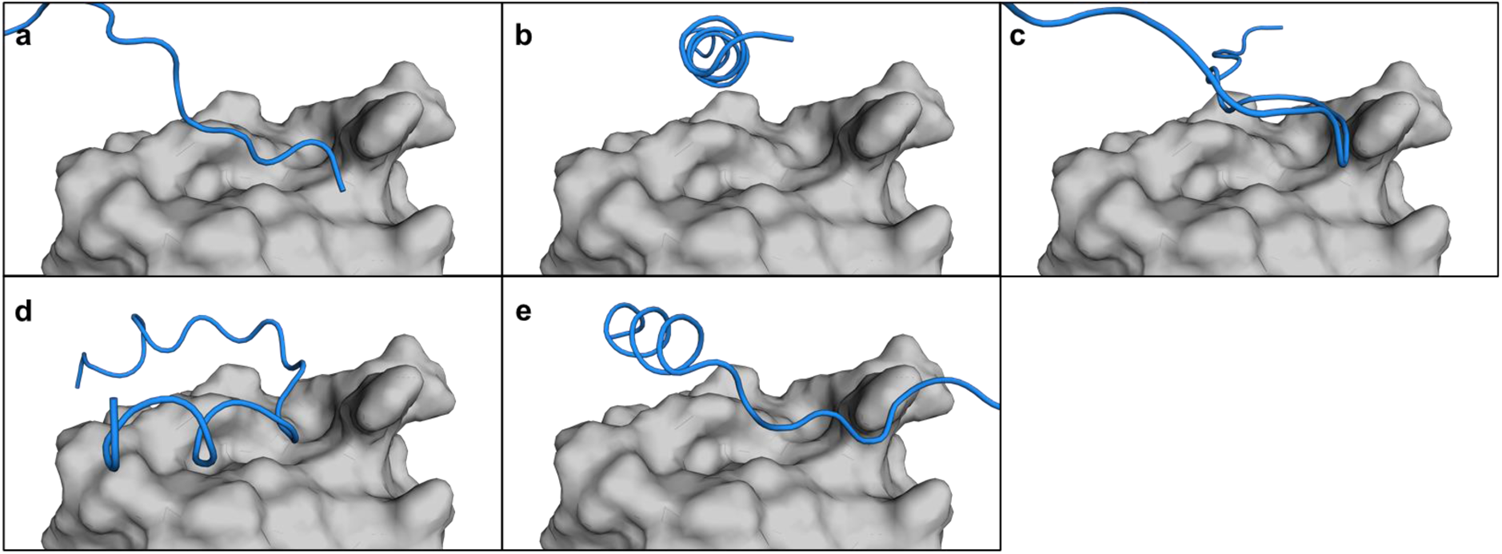
Gamma flg22 epitope permutation binding to the LRR corresponding to *BdIRI1*. **a,** Close up of the flg22 predicted binding site on LRR1 (from *BdIRI1*) with the gamma flg22 permutation #1, **b,** permutation #2, **c,** permutation #3, **d,** permutation #4, and **e,** permutation #5. LRRs are represented in grey and gamma flg22 permutation peptides are in blue. No hydrogen bonds were predicted between the LRR and any epitope permutations.

