## Supplementary material for "*Brachypodium* antifreeze protein gene products inhibit ice recrystallization, attenuate ice nucleation, and reduce immune response": File S1

File S1
All commands and scripts used
by Collin L. Juurakko, George C. diCenzo, and Virginia K. Walker


# PyMOL commands

# Basic visualization
hide lines;
show cartoon;
set ray_trace_mode, 1;
bg_color white;
set antialias, 2;
set ray_shadows, 0;
set specular, off
set ray_trace_gain, 0.0

# customize hydrogen bonds
# remove all hydrogen atoms before finding H-bonds
set dash_color, red
set dash_width, 4

# To create the closed model
# Select the residues of the LRR from the displayed sequence
# Rename selection as "LRR"
create B, LRR
# Hide everything on "LRR"
# B should be a new closed model object for the surface display without "holes" from the flg22 residues

# Remove secondary structure for flg22s
# select flg22 residues
cartoon loop

# Show side chain sticks for charged residues
#
# (script for PyMol)
# by Tom Stout, 08/05/2004
#

mstop
dss
hide all
#zoom all
#orient
show cartoon,all
color gray80,all

select pos,(resn arg+lys+his)
show sticks, (pos and !name c+n+o)
color marine,pos
disable pos
select neg,(resn glu+asp)
show sticks, (neg and !name c+n+o)
color red,neg
disable neg
set cartoon_smooth_loops,0

# this to create new flg22 object and then split it to a new chain for PISA
create flg22, (flg22)
alter flg22, chain='B'
# then select linker and original flg22 in chain A and delete
# remove secondary structure from flg22 peptide
cartoon loop, flg22

# FOR PISA WITH LRR3,4 - gamma
alter (all), chain='A'
# THEN SELECT and RENAME flg22
create flg22, (flg22)
alter flg22, chain='B'

# create colours in PymOL
set_color YellowX, [255,213,0]
set_color BlueX, [36,149,255]

# colour selected objects with new colours
color YellowX, (sele)
color BlueX, (sele)

# set surface transparency
set transparency, 0.7

# set cartoon transparency for aligned models
set cartoon_transparency, 0.3, (object)

# make everything ball-and-stick
preset.ball_and_stick(selection='all', mode=1)

# make everything sticks only
as sticks, (sele)

# To prepare multimer complexes for PISA
# select all LRRs
# where 'n' is the number of residues in one monomer
alter (sele), chain='A'
#select each LRR and add count to it
alter (sele), resi=str(int(resi)+n)
alter (sele), resi=str(int(resi)+2n)
alter (sele), resi=str(int(resi)+3n)
